# System level analyses of motor-related neural activities in larval *Drosophila*

**DOI:** 10.1101/292136

**Authors:** Youngteak Yoon, Jeonghyuk Park, Atsushi Taniguchi, Hiroshi Kohsaka, Ken Nakae, Shigenori Nonaka, Shin Ishii, Akinao Nose

## Abstract

The way in which the central nervous system (CNS) governs animal movement is complex and difficult to solve solely by the analyses of muscle movement patterns. We tackle this problem by observing the activity of a large population of neurons in the CNS of larval *Drosophila*. We focused on two major behaviors of the larvae, forward and backward locomotion, and analyzed the neuronal activity related to these behaviors during fictive locomotion that spontaneously occurs in the isolated CNS. We expressed genetically-encoded calcium indicator, GCaMP, and a nuclear marker in all neurons and used digital scanned light-sheet microscopy to record neural activities in the entire ventral nerve cord at a fast frame rate. We developed image processing tools that automatically detect the cell position based on the nuclear staining and allocate the activity signals to each detected cell. We also applied a machine learning-based method that we developed recently to assign motor status in each time frame. Based on these methods, we find cells whose activity is biased to forward versus backward locomotion and vice versa. In particular, we identified a group of neurons near the boundary of subesophageal zone (SEZ) and thoracic neuromeres, which are strongly active during an early phase of backward but not forward fictive locomotion. Our experimental procedure and computational pipeline enable systematic identification of neurons to show characteristic motor activities in larval *Drosophila*.

## Introduction

An animal responds to changes in the external and internal environment by choosing the appropriate behavior among the animal’s repertoire^1–3^. Each behavior of an animal is generated by a series of muscle contractions in different parts of the body, which is induced by spatio-temporally coordinated activation of motor neurons innervating these muscles. The motor neuron activity in turn is regulated by networks of interneurons in the CNS. How specific motor activity emerges in neuronal circuits remains poorly understood. Previous studies in animals with relatively small nervous systems suggested that dynamics of neural activity in a population of neurons determines the selection and execution of a specific behavior^3,4^. It would therefore be crucial to study the activity of neurons at the system level to understand how a specific sequence of motor activity emerges in the central circuits. Due to technical difficulties, system-wide activity recording has previously been limited to animals with fewer than hundreds of neurons such as leeches and worms^4,5^. However, with the development of fluorescent probes and measuring instruments over the last decade, it is becoming possible to measure the activity of neural populations in animals with a larger CNS.^6–8^.

*Drosophila* larvae provide an excellent model system for studies of neural circuit mechanisms underlying motor regulation. During forward locomotion, the larvae contract muscles in the abdominal segments of the body wall from the posterior (A8) to anterior (A1)^9,10^. During backward locomotion, direction of the propagation of muscle contraction is reversed. When the larvae change direction, they execute the turning behavior, in which muscles in one side of the anterior body is contracted. The larval CNS consists of the brain, subesophageal zone (SEZ) and ventral nerve cord (VNC) and, contains ~ 10,000 neurons. The VNC is further divided into thoracic and abdominal neuromeres. Since motor neurons in each neuromere innervate muscles in the corresponding segment of the body wall^11^, the patterns of neural activity of motor neurons in the VNC reflect those of muscle contraction during larval behaviors. Therefore, it is possible to infer the actual larval behavior from the activity pattern of the motor neurons in the CNS^9^. Furthermore, neural activities corresponding to forward and backward locomotion and turning can occur in isolated CNS^12^ (fictive locomotion), providing a good opportunity to study activity dynamics underlying motor generation without being disturbed by the movement of the animal. Taking advantage of strong genetic tools and recently developed connectomics, previous studies identified segmental interneurons and circuits present in each abdominal neuromere that regulate motor patterns, such as speed, intersegmental coordination, left-right coordination and sequential recruitment of motor pools^13–18^. Previous studies also identified sensory-motor interneurons which elicit specific behaviors, such as rolling and backward locomotion, in response to sensory stimuli^19,20^. However, these interneurons likely represent only a small fraction of interneurons recruited during the execution of larval behaviors, since a large number of interneurons are yet to be characterized.

In this study, we aimed to study neural activities related to larval locomotion at the system level. We used a digital scanned light-sheet microscopy combined with calcium imaging to record neural activity in a large number of neurons in the VNC, SEZ, and part of the brain during fictive forward and backward locomotion. We developed image processing tools that identify cells in the calcium imaging data and extract activity signals in the cells. We also applied a machine learning-based method that we developed recently^21^ to identify the time windows in which forward and backward locomotion occurs. Taken together, these methods enabled systematic and objective analyses of the activity of a neural population in this system.

## Results

### Automated cell detection and signal extraction

We used a digital scanned light-sheet microscopy ezDSLM^22^ to record neuronal activity in isolated CNS preparations from first instar larvae. We expressed a genetically encoded Ca^2+^ sensor GCaMP6f for activity recording and nuclear marker mCherry.nls for cell detection, pan-neuronally with the *elav-Gal4* driver. To archive both high temporal resolution required for activity recording and high spatial resolution required for cell detection, we performed imaging under two different conditions: high temporal resolution (0.6s/volume) but low spatial resolution (along the z-axis, z-stacks of 32 planes acquired with 2.485*µ*m step size) imaging for GCaMP6f signal (called a dynamics image, Fig. 1a, b) and high spatial resolution (z-stacks of ~ 1000 planes acquired with 0.146*µ*m step size) imaging for GCaMP6f signals (an intermediate image, Fig. 1a’, c) and mCherry.nls signals (a reference image, Fig. 1a”, d). The dynamics image was registered with the reference image via the intermediate image, which includes the GCaMP6f signal captured simultaneously with the reference image at the same spatial resolution.

**Figure 1.**
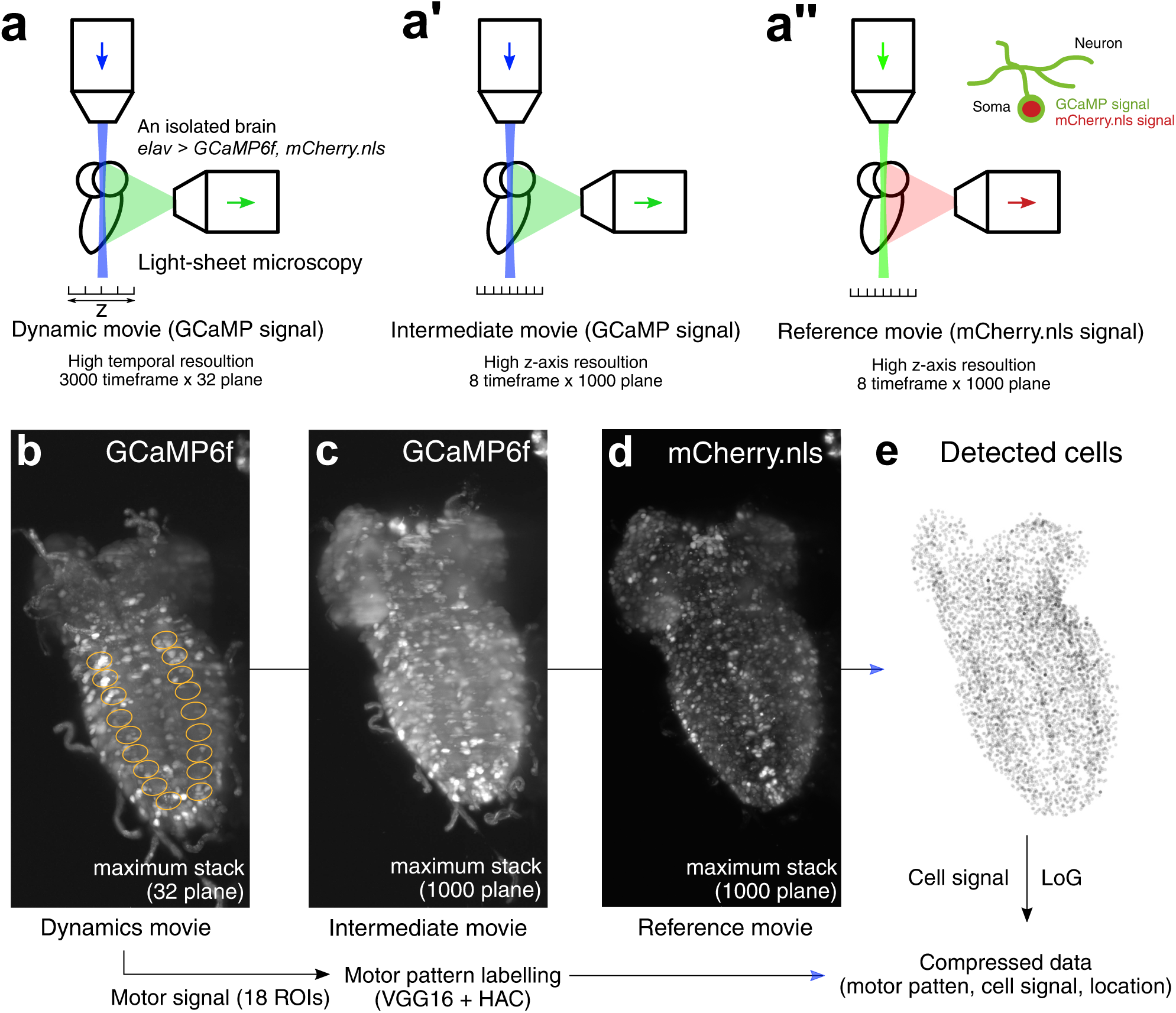
Outline of automated cell detection. (a-a”) Diagrams showing imaging protocols. (b-e) Actual images. Calcium signals and nuclear locations were imaged in an isolated CNS from an *elav>GCaMP6f, mCherry.nls* 1st instar larva. First, activity signals (GCaMP) were imaged with relatively few focal planes (~30) at a fast frame rate (~0.6s/volume) for ~30 minute (dynamics image; a, b). Then, GCaMP and nuclear (mCherry.nls) signals were simultaneously imaged at a high z-axis resolution to obtain an intermediate image (a’, c) and reference image (a”, d), respectively. (e) Cells detected by computational analyses. By aligning the dynamics image with the reference image via the intermediate image, activity signals are allocated to individual cells. By analyzing motor signals in the 18 ROIs in the dyanamics movie (circles in b), motor status in each time frame was also determined. See Methods for detail.

We imaged neuronal activity in the entire VNC and part of the brain (dorsal brain regions were excluded from the imaging to increase temporal resolution), which together include thousands of neurons. We detected the cells by applying Laplacian of Gaussian (LoG) and Hessian operation to the reference image that visualizes the cell nuclei (see Methods for detail). Comparison of the locations of part of the nuclei thus assigned (Fig. 1e) with those annotated by human inspection (127 cells) indicated that the automated detection method yields 86.2% hit. Activity data were then obtained from the dynamics image by extracting GCaMP6f fluorescence signals in the detected cells.

### Classification of motor states

In characterizing the activity data of ~ 5,000 neurons acquired, we first determined the time windows in which activities reflecting specific larval behaviors such as forward and backward locomotion occur by adapting the automated classification of motor states we developed in another study^21^. The method utilized convolutional neural network (CNN) and cluster analysis and successfully classified motor patterns in various conditions. The idea behind the use of CNN is that images representing motor activity pattern are similar to those of handwritten digits and thus would be effectively classified by CNN. As in the previous study, we extracted activity signals in the regions of interest (ROIs) representing 18 hemineuromeres in the dynamics image (Fig. 1b) and compressed it to a 9-d time series by adopting the maximum value in each neuromere. The 9-d time series is divided into windows and transformed to 8-bit x-t images in which the x-axis represents the activity of the 9 neuromeres at each time point (see Methods for detail) We then extracted features of the images from the 3rd layer of the 4th convolutional block (Conv4 3) of the CNN model VGG16, which the previous study showed to give best performance among the CNN layers, by global average pooling and clustered the features by hierarchical agglomerative clustering (HAC, Ward’s method). By post-hoc human labeling of the clusters, we could define motor patterns corresponding to a forward wave (FW), backward wave (BW) and left-right symmetric or asymmetric activation in anterior neuromeres (AT) and quiescence state (QS) (Fig. 2a, b), as in the previous study. As was observed in the previous study, AT often occurs just before BW and together with BW appears to constitute the fictive backward locomotion. Unlike in the previous study, the motor pattern corresponding to synchronous and bilateral activation in posterior-most neuromeres (called PT), which often occurs prior to FW, was not detected in this study and appears to be merged with FW, likely due to difference in the target cell populations and/or imaging conditions. By applying the classification to the imaging data, we assigned the time windows of each motor pattern and thus were able to investigate the activity of the ~ 5,000 neurons during each motor pattern (Fig. 2c). Overall inspection of the activity pattern with the assigned motor states confirmed the validity of the classification (e.g., activity propagation from the posterior to anterior CNS coincides with FW). As an independent method for detecting and distinguishing motor patterns, we also performed principal component analysis (PCA) on the time derivatives of the cellular signals. We found that the first three principal components accounted for 51% of the variations in the population neural activity. For each PC, a corresponding time series (temporal PC, TPC) was calculated and plotted in the three dimensional PC space (Fig. 2c, d). When time windows of motor states AT, BW, and FW assigned by the automated motor classification above were plotted in the TPC space, they were found to correspond to three distinct neural state trajectories (Fig. 2d).

**Figure 2.**
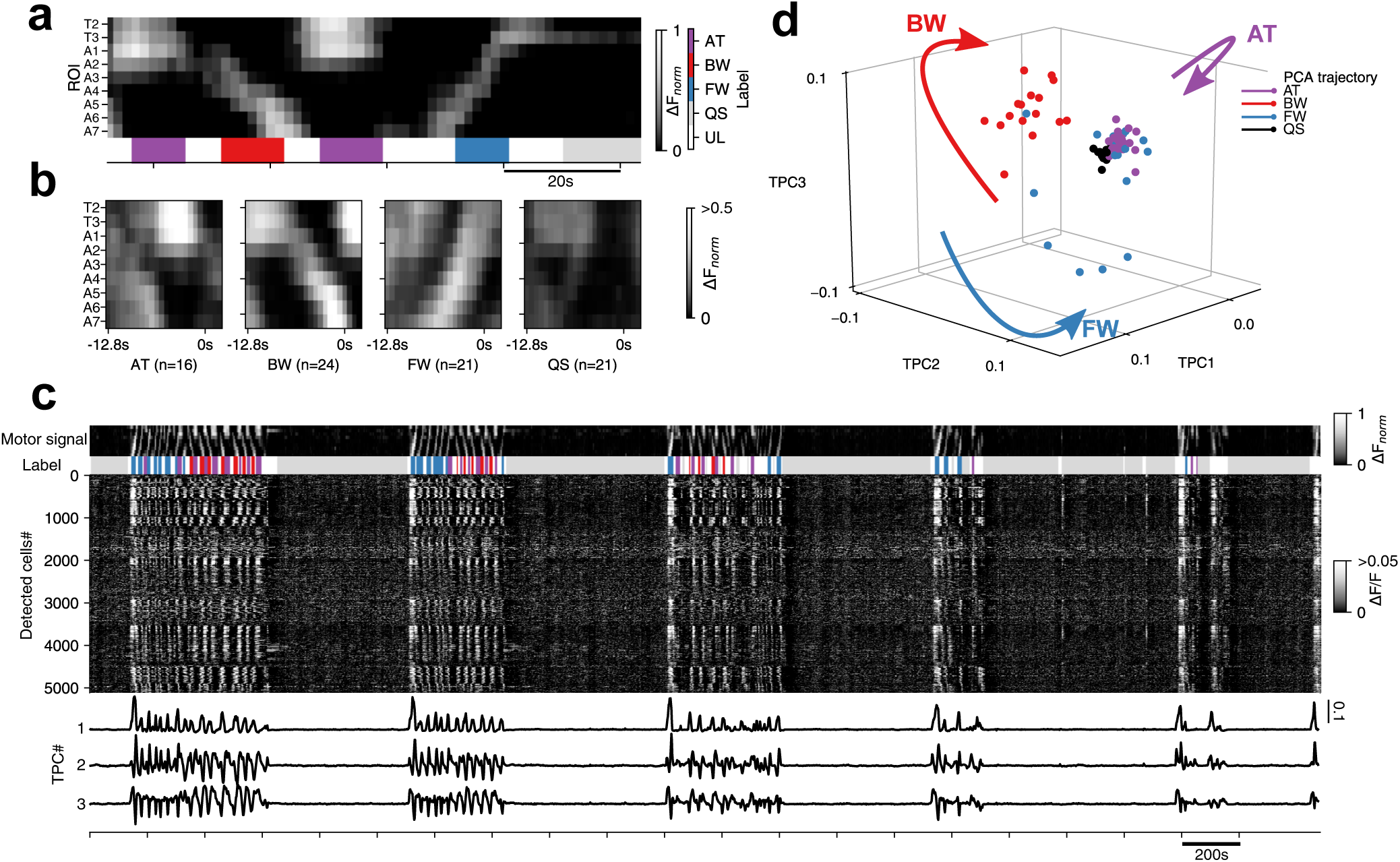
Motor patterns and neuronal activities. (a) A sample time interval showing activities in 9 regions of interest (ROI 1-9, from the anterior to posterior neuromere) and assigned motor status represented in different colors as indicated in the color-code bar. AT, anterior burst; BW, backward wave; FW, forward wave; QS, quiescence state; UL, unlabeled. (b) Averaged activities in the 9 ROIs during each fictive motor pattern. (c) Activities of the entire recorded cells (middle) and temporal principal component (TPC, bottom), aligned with motor activities in the 9 ROIs and assigned motor status (top). Cells were numbered along the anterior-posterior axis with cell 1 being the most anterior. (d) Dynamics of the whole population neuronal activities visualized in 3D PCA space. Trajectories are colored for motor patterns with circles indicating the end point of each motor pattern. Arrows indicate the direction of the trajectories.

### Candidate backward/forward wave biased cells

Having assigned the time windows when forward or backward fictive locomotion occurs, we next searched for cells whose activity is correlated with the motor patterns. To identify candidates of backward/forward wave biased cells, we profiled the activity of the cells. The maximum value of ∆F/F during the period of BW, FW and AT are calculated in each cell (called BW_*max*_, FW_*max*_ and AT_*max*_, respectively). The time periods used in the calculation were extended to include a few seconds prior to the event (2.4s for FW and BW and 1.2s for AT), since some of the neurons involved in motor regulation may be active during these periods. We defined motor pattern-biased cells according to the following criteria: BW-biased cells, BW_*max*_ > 0.05, FW_*max*_ < 0.055 and AT_*max*_ < 0.05; FW-biased cells, BW_*max*_ < 0.05, FW_*max*_ > 0.08 and AT_*max*_ < 0.09. These parameters were determined by manual fine-tuning so that *<* 200 cells would be identified under each criterion. The profiling identified ~ 150 biased cells for each motor pattern (Fig. 3a). We noticed that many of the FW-biased cells are distributed in the dorsal VNC and likely include motor neurons.

**Figure 3.**
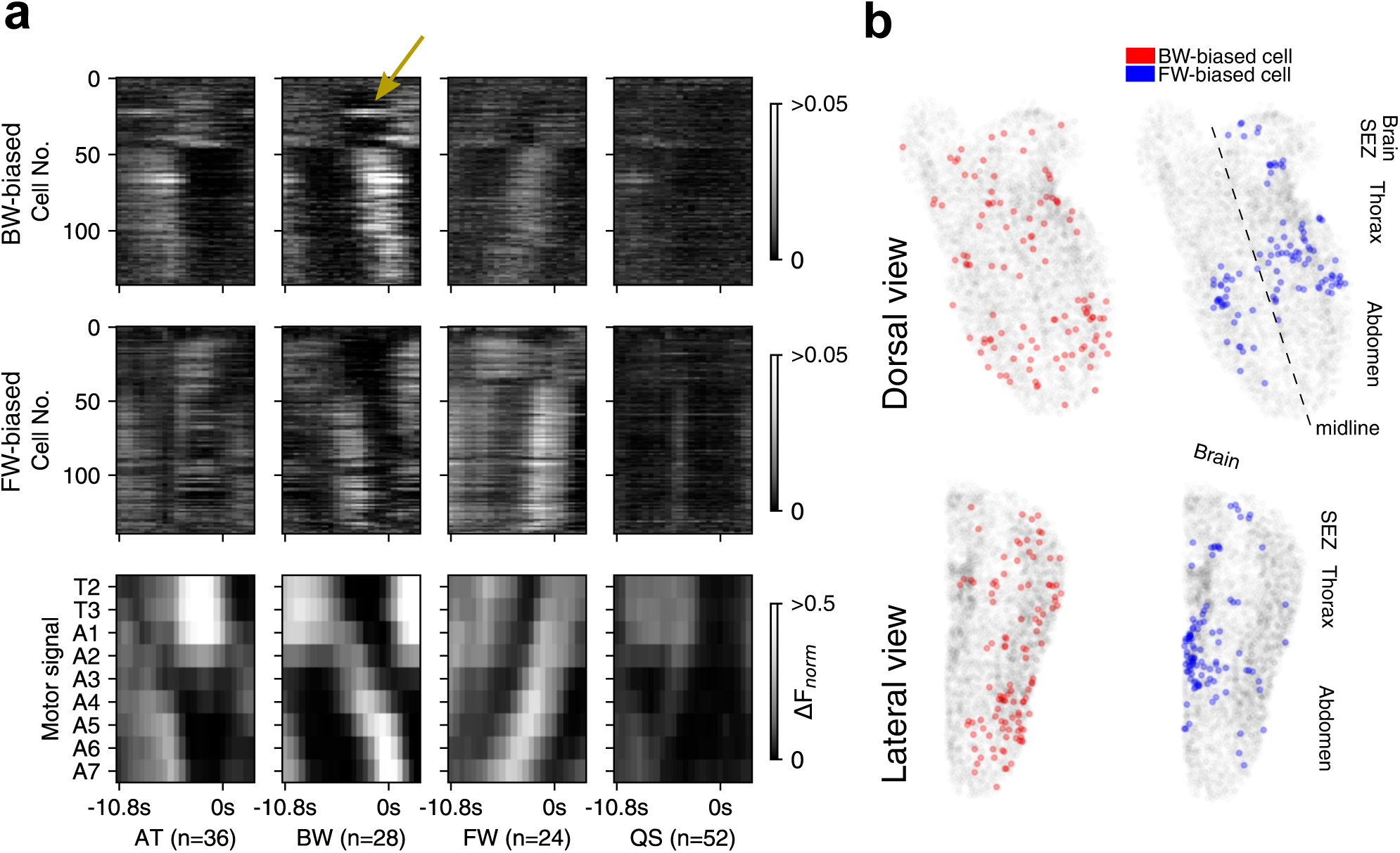
Candidate backward/forward wave biased cells. (a) Average activities of candidate BW-biased cells (top) and FW-biased cells (middle) during each motor patterns. Motor activities are also shown (bottom panel). Cells are sorted from the anterior (top) to posterior (bottom). Arrow indicates a group of neurons which show the earliest activities during BW and are further examined in Figure 4. (b) Distribution of BW-biased cells (left, red) and FW-biased cells (right, blue). A dorsal view (top) and lateral view (bottom) are presented.

Among the candidate FW and BW-biased neurons, we focused on neighboring four BW-biased cells (B-21,22,23,25), which are activated the earliest among the BW-biased cells (arrow in Fig. 3a). As shown in Fig. 4a, b, these cells are strongly active during an early phase of BW but only weakly during FW. They are located near the ventral midline and near the boundary of SEZ and thoracic neuromeres (Fig. 4c). This CNS region has previously suggested to include neuronal circuits that initiate backward locomotion^20^ (see Discussion). The location and timing of their activity make these neurons good candidates for interneurons regulating backward locomotion.

**Figure 4.**
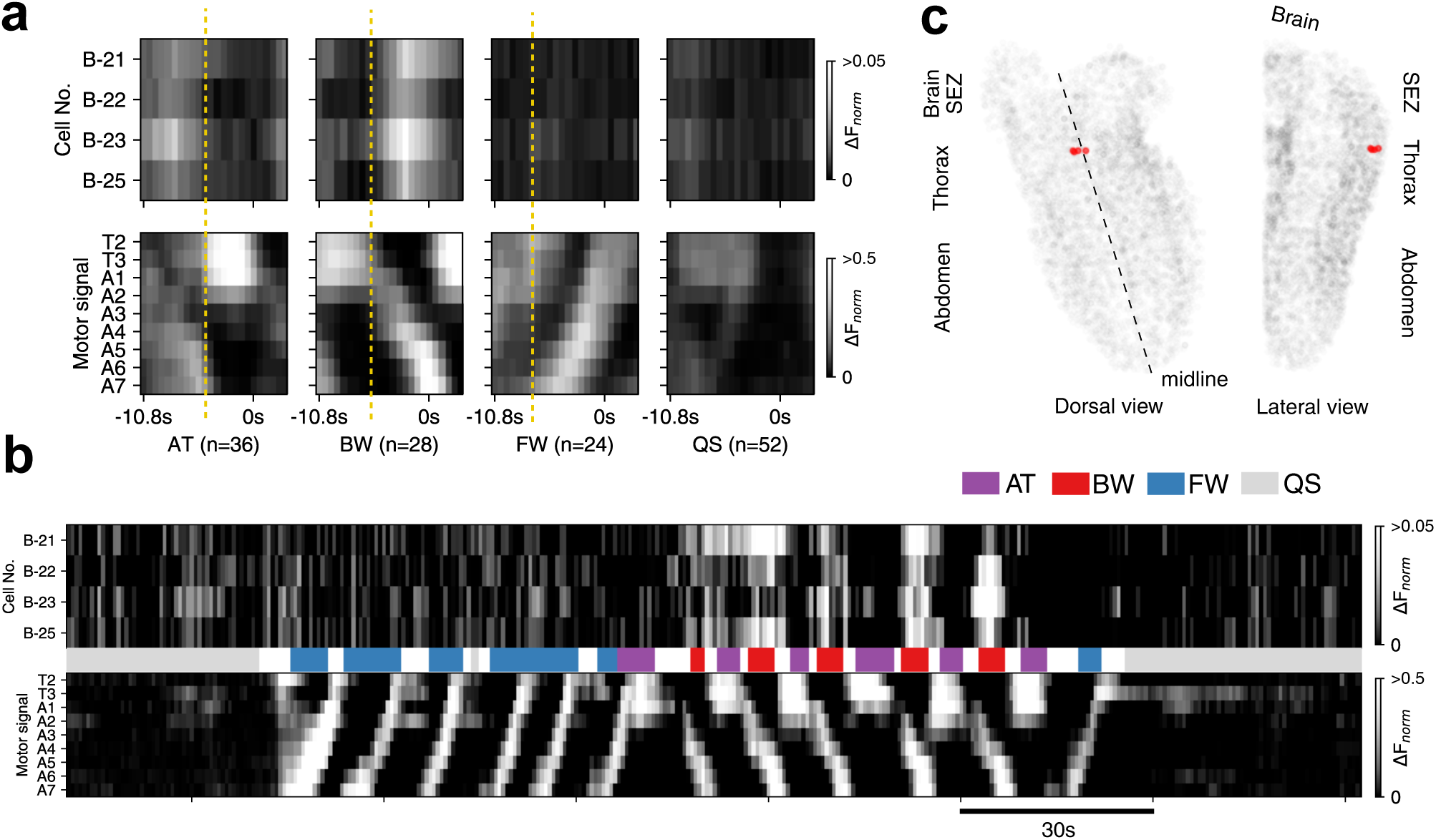
Cells preferentially active during an early phase of backward waves. (a) The average activities of BW-biased cells B-21, 22, 23, 25 (top) during each motor pattern, aligned with the motor activities (bottom). Dotted lines indicate the initiation of each motor pattern. (b) A sample time interval showing activities of the four cells (top), aligned with the motor status (middle, color coded) and motor activities (bottom). (c) Position of the four cells. They are located at the boundary between the SEZ and thorax.

## Discussion

We recently introduced the use of machine learning-based computational methods to classify motor states in larval Drosophila^21^. In the study, spinning-disc microscopy was used to image neuronal activity in the dorsal VNC containing the neuropile region in the 3rd instar larvae. CNN combined with cluster analyses successfully identified the motor states in the calcium imaging data, in which GCaMP6f was expressed in all or subsets of neurons (*eve*-expressing motor neurons and *5-HT2A-GAL4* expressing and *trh-GAL4* expressing interneurons). In the present study, we applied light-sheet microscopy to a smaller 1st instar CNS to image the entire VNC. The same CNN layer and cluster analysis was successfully used to detect the motor states in this preparation. Furthermore, by co-visualizing the nuclei, we allocated the activity signals to individual cells. While we could record the activity of a majority of neurons in the SEZ and VNC during fictive locomotion, a large part of the brain was excluded from the analyses to increase the temporal resolution. However, previous work suggested that the larvae can crawl in the absence of the brain and SEZ^23^. Furthermore, previous ablation experiments showed that fictive forward and backward locomotion can occur even when the brain and SEZ are ablated^12^. Thus, the VNC, on which we focused our imaging, should include self-contained circuits that can autonomously produce the motor activities.

Having determined the motor state in each time frame, we next searched for and found neurons whose activity is biased toward forward or backward fictive locomotion. Both FW-biased and BW-biased neurons were found to exist in a largely left-right symmetrical manner and throughout the anterior-posterior axis of the VNC. It suggests that the circuits regulating forward and backward locomotion are scattered in the VNC. This is consistent with the previous ablation experiments which suggested that any segment from A2/3 to A8/9 is able to initiate both forward and backward waves^12^. Identified biased neurons included a cluster of neurons near the midline at the boundary between the SEZ and thoracic segments, which are among the earliest to be activated during BW. The location of these cells overlap with the region that contains downstream circuits of Wave neurons, command-like neurons for backward locomotion^20^. Wave neurons are segmentally repeated interneurons present in abdominal segments but differ in function depending on their location along the anterior-posterior axis: anterior Wave neurons function as command neurons that elicit backward locomotion in response to touch stimuli in the head whereas posterior Wave neurons elicit forward locomotion in response to touch stimuli in the tail. While Wave neurons are activated by the sensory stimuli but not during spontaneously occurring fictive backward locomotion, the downstream neurons likely involve those that are specifically active during backward locomotion and actuate the motor program. The BW-biased neurons identified in this study may include such neurons. Previous studies also used sparse Gal4 lines to identify interneurons which are active during forward and/or backward locomotion and implicated in the regulation of larval locomotion^13,14,16–18,21^. These studies identified several interneurons whose activity are biased to FW or BW, including FW-biased A27h^17^ and CLI1^18^ neurons and BW-biased Leta neurons^21^. Since many FW- or BW-biased neurons were identified to be present in the VNC in the present study, it was difficult to determine whether these include the previously identified FW- or BW-biased neurons. Further characterization of each of the candidate FW- and BW-biased neurons (e.g., with molecular markers) is required to address this issue.

While this work was in progress, a similar study on *Drosophila* larval functional imaging was published by Lemon et al.^24^. The authors used state-of-the-art multi-view light-sheet microscopy with one or two-photon excitation to achieve superior temporal and spatial resolution in the 1st and 3rd instar CNS. In the study by Lemon et al.^24^, GCaMP alone was expressed in neurons and thus functional unit of the statistical analyses of neuronal activity was voxel but not cells. Instead, in this study, a nuclear marker was co-expressed to allow activity profiling at the cell-level. Our study also identified BW-biased neurons that are not described by Lemon et al.^24^. Difficulty common to our study and Lemon et al.^24^ is that since all neurons are visualized, it is difficult to know the identity of the neurons that show characteristic activity patterns. Our previous study showed that when GCaMP is expressed in fewer than ~ 10 cells in each neuromere, correlation analyses may be used to reveal the outline of the neuron showing specific activity^21^. Thus, by applying the functional imaging technique such as the one described in this study to a large number of relatively sparse Gal4 lines may enable system level analyses of motor activity while retaining the ability to identify the neurons of interest. Once candidate neurons are identified, their roles may be studied by the use of optogenetics. Furthermore, by combining functional imaging with optogenetical perturbations, system-level analyses of the dynamics of the motor circuits would be possible in the future.

## Methods

### Strains

We used the following fly strains in this study: *elav-Gal4* (Davis et al., 1997), *UAS-GCaMP6f* (Bloomington Drosophila Stock Center (BDSC) stock No. 42747), and *UAS-mCherry.nls* (BDSC stock No. 38424).

### Calcium imaging

We isolated the larval CNS (central nervous system) from the first instar larvae (21 ~ 30 hours after egg laying) to prevent muscle contraction from interfering calcium imaging. The isolated CNS was mounted on the MAS (Matsunami adhesive silane)-coated slide glass (Matsunami, Osaka, Japan) and submerged in the TES buffer (TES 5 mM, NaCl 135 mM, KCl 5 mM, CaCl_2_ 2 mM, MgCl_2_ 4mM, Sucrose 36 mM). To perform 4-dimensional imaging, we used ezDSLM^22^, a digital scanned light-sheet microscopy (DSLM). By using the ezDSLM system, we can record calcium signal from multiple focal planes (~ 30) at a rate of 0.6 s/volume. To obtain a single data set, firstly, we recorded neural dynamics of the larval CNS (called dynamics image). In the dynamics image, GCaMP6f fluorescence from fewer (32 planes), and sparser (2.485 *µ*m*/*plane) focal planes are acquired so as to achieve high temporal resolution (0.6 s/volume). After recording the dynamics image, we scanned GCaMP6f and mCherry.nls (nuclear-localization signal) fluorescence in more (1000 planes) and denser (0.146 *µ*m*/*plane) focal planes to detect neuron position more precisely. The GCaMP signal (called intermediate image) and mCherry.nls signal (called reference image) in the second scanning were taken in an interleaved manner, so that there is no need for spatial alignment between the intermediate image and the reference image. The parameters used for the recordings are summarized in the following table.

**Table 1.** Parameters of movie acquisition

| Parameters | Dynamics | Intermediate | Reference |
| --- | --- | --- | --- |
| Dimension (normal) | 32 planes | 1000 planes |  |
| Dimension (focal) | $2048 \times 1024 \text{ px}^2$ | | |
| Interval (normal) | $2.485 \text{ }\mu\text{m/px}$ | $0.146 \text{ }\mu\text{m/px}$ | |
| Interval (focal) | $0.146 \times 0.146 \text{ }\mu\text{m}^2/\text{px}^2$ | | |
| Temporal resolution | 0.638 s | - |  |
| Laser wavelength | 488 nm |  | 561 nm |

### Registration and preprocessing

We developed a registration method to correct the translation of the sample in the movies (https://github.com/Jeonghyuk/Canal-Yoon). Since our data showed no obvious drift in the z direction and rotation, we corrected drift in the x and y axes. First we stacked the data in the z direction. To obtain the offset in the x and y directions, we created lattice-like points in the image at frame t as feature points and generated pieces of images (256 × 256 pixel) around the feature points. By minimizing the normalized cross-correlation between the pieces at frame t and the whole image at frame t+1, we obtained the optimal offset in the x and y directions to correct the drift. By repeating this process, we corrected all drifts in the movies.

To find the area of the CNS and the neuropil of the samples, we used the reference images in which mCherry proteins visualize *the nuclei* of the all neurons in the CNS and the intermediate images, in which GCaMP6f proteins visualize the *whole cytoplasms* of the neurons. We estimated the boundary region of the CNS using the intermediate images and the Sobel-Feldman operator (see below). After detecting the CNS region, we determined the neuropil region by subtracting the bright region in the reference images, which highlight cell nuclei and thus the cortex region, from the CNS region. We excluded the neuropil region in the cell detection process to suppress false positive detection of somas in the neuropil.

The Sobel-Feldman operator^25,26^ creates an image which emphasizes the edges in the 2-dimensional input image. The operator uses 3 × 3 kernels, *G*_*x*_ and *G*_*y*_:

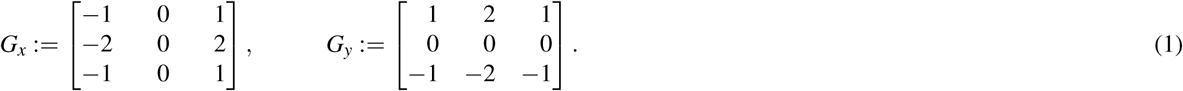

Applying the above matrices for a 2-dimensional image *f*, we obtained images *f*_*x*_, *f*_*y*_ which are approximations of the gradient of the input image along with the *x*- and *y*-axis respectively:

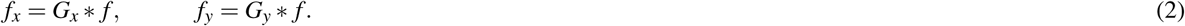

Since the gradient magnitude 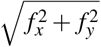 of the images has a high value in the boundary region, the border of the sample could be estimated using the Sobel-Feldman operator.

### Cell detection

The reference images (images of the nuclei) were used to perform cell segmentation. The movies were taken in an isotropic manner in xyz axes (0.146 *µ*m*/*px) by the settings of the recording. Using the CNS mask and the neuropil mask described above, we excluded voxels which are outside of the CNS area or inside of the neuropil area to focus on the cortex region, which includes cell bodies. Within the cortex region, we first detected all local maxima including the vertices and edges by applying LoG (Laplacian of Gaussian), a method commonly used for local maxima detection in an image ^27^ as follows. For the 3D image *f*: ℝ^3^ → ℝ, the LoG {*f*} generates a 3D image expressed as the following definition:

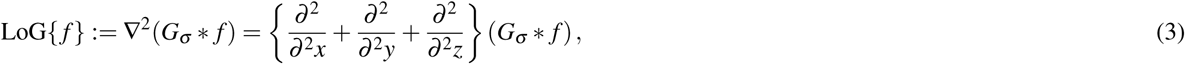

which is the Laplacian filter applied after blurring the image. *G*_σ_ is a Gaussian probability density function with zero mean and standard deviation *σ*. Smoothing with *G*_σ_ reduces noises below *σ* pixels.

LoG not only detects the center of the cell that we focus on but also the points corresponding to the edges where the change of pixel value is large. To exclude the points corresponding to the edges, we used an index^28^ based on Hessian *H*(*G*_*σ*_ * *f*) of the 2D image defined as follows,

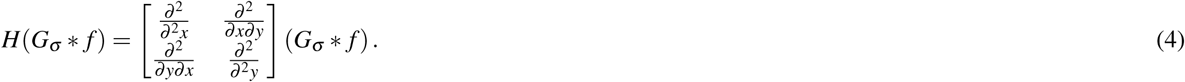

The principal curvature of the points is the ratio *r* = *α*/*β*, where the first and second eigenvalues of the Hessian is *α* and *β*, respectively. If the principal curvature satisfies *r* ≪ 1 or *r* ≫ 1, the point corresponds to an edge. To compute the principal curvature, we use the following relational expression, in which we don’t need to calculate the eigenvalues:

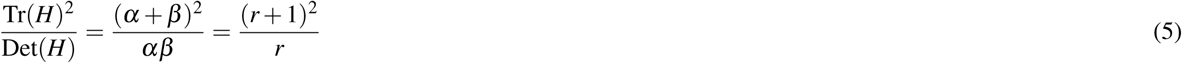

Fluorescence time series of the cells are extracted from the dynamics image, using the center of cells detected as described above. To cancel out the bleaching effects, we normalized the signals by dividing the original data by a baseline. In obtaining the baseline, we first calculated a local minimum value in each time window (from t-7 to t+7). Then the Gaussian blur with *σ* = 7 frames was applied at each frame to obtain the baseline.

### Motor patten classification

Motor pattern classification was performed as previously described^21^. We measured changes in fluorescence intensity in 18 ROIs, one for each hemisegment and in a dorsal neuropile region containing neurites of motor neurons (nine left and nine right, from T2 to A7). The baseline (F_*base*_(t)) was calculated as the mean fluorescence signal from t 16 to t within each ROI. The changes in fluorescence (∆F/F) were defined as (F(t)/F_*base*_(t) 1)H(F(t)/F_*base*_(t) 1) where H(x) is the Heaviside function. For image conversion, ∆F/F is scaled to between 0 to 255 (8 bit integer) using the minmax transform. 9-d time series were calculated using the maximum value among the left 9 ROIs and right 9 ROIs. We defined the window size as 8 time frames (from t-2 to t+5) so that each window has 9-d time series with 8 time frames. Each window was transformed to a 72 pixel x 72 pixel RGB image where the RGB channels have the same value. The features of the images were extracted using global average pooling from the Conv4_3 intermediate layer of ImageNet pre-trained VGG-16 model^29^ and clustered by HAC using Ward’s method. Labeling was performed comparing average images of the first ~24 clusters. Images showing high intensity in T2-A4 neuromeres are categorized as AT. Those showing angled intensity flow are categorized as FW or BW depending on the direction of the flow. Those showing nearly no intensity change are categorized as QS. PCA was performed on the time derivatives of ∆F/F of all cells (5117 cells).

## Acknowledgements

We are grateful to Bloomington Drosophila stock center and Kyoto stock center. This work was supported by the Advanced Leading Graduate Course for Photon Science (Y.Y. and J.P.) and a MEXT/JSPS KAKENHI grant (15H04255, 16H06280, 17K19439, 17H05554 and 22115002)

## Author contributions statement

Y.Y., H.K., S.I., S.N. and A.N. conceived and designed the experiments. Y.Y., A.T. performed the experiments. Y.Y., J.P., K.N. analyzed the data. H.K. contributed reagents. Y.Y., J.P., H.K. and A.N. wrote the main manuscript. All authors reviewed the manuscript.

## Additional information

**Competing financial interests** The authors declare no competing financial interests.

